# Discovering Condition-specific Cell Populations via Integrative Clustering of Single-cell Data

**DOI:** 10.1101/2025.10.03.680413

**Authors:** Rekha Mudappathi, Bhardwaj Vaishali, Li Liu

## Abstract

We present <u>IN</u>tegrative CLustering Of Single cElls (INCLOSE), a novel computational method that integrates single-cell omics data and sample metadata to identify cell populations. INCLOSE analysis of CITE-seq data of acute myeloid leukemia and healthy samples uncovered cell populations exclusively found in the leukemia samples or the healthy samples. These condition-specific cell populations strongly suggested that immune suppression in tumor microenvironment plays a pivotal role in driving tumor progression.

## Main text

Cell populations in a tissue continuously adapt to environmental stimuli, changes in health status, and other intrinsic and extrinsic factors ^1^. During the process, various cell populations may emerge, expand, contract, or disappear entirely. Identifying these distinct cell groups is essential for unraveling cellular heterogeneity and dynamics.

An important aspect of single cell omic data analysis is the identification of cell populations. Typically, the process involves pooling cells from all sequenced samples, conducting a clustering analysis, and subsequently visualizing the cluster distributions respective to experimental conditions, such as by treatment or disease groups ^2^. This conventional method has a limited power to uncover condition-specific cell populations that tend to be underrepresented across all samples and merge into larger or somewhat similar clusters ^3^. An alternative approach is to cluster cells from different subsets of samples separately, but it is difficult to find the correspondence between clusters inferred from independent clustering analyses. To address this challenge, we have developed <u>IN</u>tegrative CLustering Of Single cElls (INCLOSE), a novel computational method that incorporates sample metadata in clustering analysis. INCLOSE is versatile, capable of integrating single-omic or mulit-omic data at the single cell level with sample metadata that may contain a single variable, such as a disease group, or include additional covariates.

We use an existing cellular indexing of transcriptomes and epitopes (CITE-seq) data set to demonstrate the INCLOSE algorithm. CITE-seq allows for simultaneous quantification of a multitude of cell surface proteins and the transcriptome in the same cell ^4^. Knorr *et al*. conducted a CITE-seq experiment on eight individuals newly diagnosed with accurate myeloid leukemia (AML) and three age-matched normal donors ^5^. A total of 131 surface proteins and 22,998 gene transcripts were measured. Using the Seurat ^6^ and MAESTRO ^7^ pipelines (**Online Methods**), we identified ten cell clusters, among which myeloid cells (MCs) expressing CD11b(+)CD33(+)HLA-DR(low) ^8^ consisted of 2,543 and 4,234 cells in the healthy group and the AML group, respectively (**Supplementary Fig. 1**). Because MCs are highly heterogeneous and different subtypes of MCs in tumor microenvironment have been associated with AML progression ^9^, we applied INCLOSE method to further identify condition-specific MC subpopulations.

INCLOSE started with estimating pairwise correlations between cells (**Fig. 1A**). It computed two correlation matrices, one using the top 2,000 highly variable genes among MCs, and the other using the top 15 highly variable surface proteins. A third correlation matrix was constructed using the metadata by assigning values of 1 to cells from the same sample group (healthy or AML) and 0 to cells from different groups. The three correlation matrices were then fused into a cell-cell similarity matrix through weighted multiplication where the weights controlled the relative contribution of each data modality to cell-cell similarities. INCLOSE also applied Gaussian smoothing to reduce noise caused by minor variations in correlation coefficients. The clustering process initiated with identifying a seed cell that exhibited the highest overall similarity. Other cells were then ranked based on their similarity to this seed cell and progressively grouped into a cluster until the within-cluster similarity stopped increasing ^10^. The seeding and expanding steps were repeated for the remaining cells until no cluster with more than 5 cells could be formed (**Fig. 1B**). To assess the quality of these clusters, INCLOSE calculated within-cluster similarity and between-cluster distance and combined them into a segregation score (**Online Methods**).

**Figure 1.**
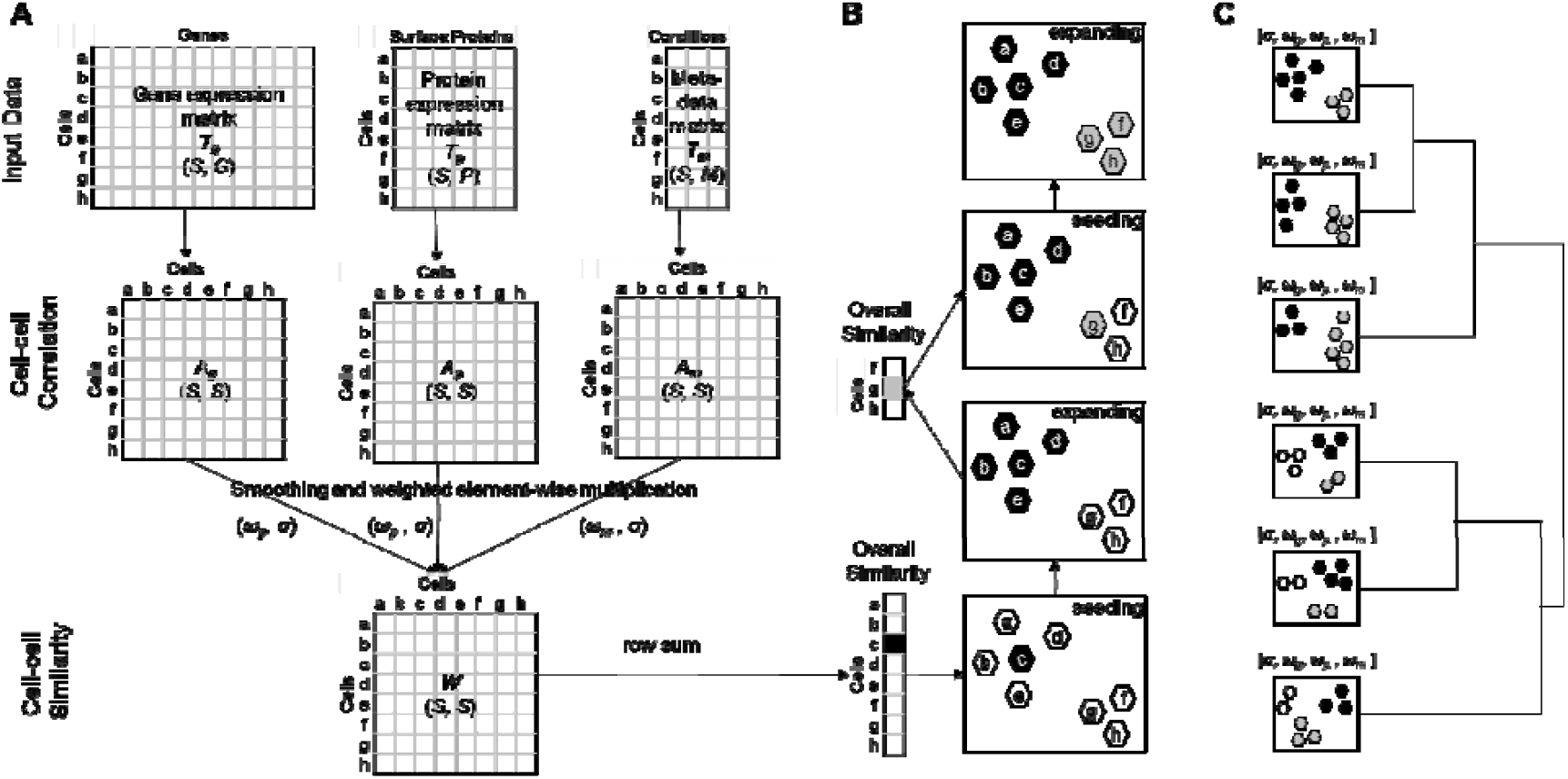
Schematic illustration of the INCLOSE algorithm. (**A**) S cells are characterized by expression of G genes and P surface proteins and M meta features. Pairwise correlation between cells is calculated based on each data modality, which are subsequently combined into a cell-cell similarity matrix tuned by weights (ω) and smoothing parameter (σ). (**B**) The clustering process involves a series of seeding and expanding steps. At each iteration, the cell with the highest overall similarity serves as the seed. (**C**) Various combinations of tuning parameter values are tested. The clustering results are organized into a hierarchical tree based on J-score that quantifies alignments between two clustering results.

Using different weights and smoothing parameter leads to different clustering results. The optimal combination of these tuning parameters was not known *in priori*. INCLOSE addressed this issue by iterating through a series of parameter values and performing clustering analysis for each possible combination. To facilitate exploration and visualization of these clustering results, INCLOSE aligned them via bidirectional set matching, calculated the alignment J-score ^11^, and organized them into a dendrogram (**Fig. 1C**).

We performed INCLOSE analysis to cluster the MCs from the AML study, exploring a total of 177 distinct combinations of tuning parameters. The segregation scores of these clustering results ranged from 0.0008 to 0.935 (**Supplementary Fig. 2**). The 45 clustering results with segregation score >0.9 were distributed across four clades in the dendrogram (**Fig. 2A**). Within each of these four clades, the clustering results exhibited a high degree of similarity, with the number of clusters ranging from six to eight, and J-scores ranging from 0.779 to 0.999. Notably, all these clustering results revealed condition-specific clusters (**Supplementary Fig. 3**).

**Figure 2.**
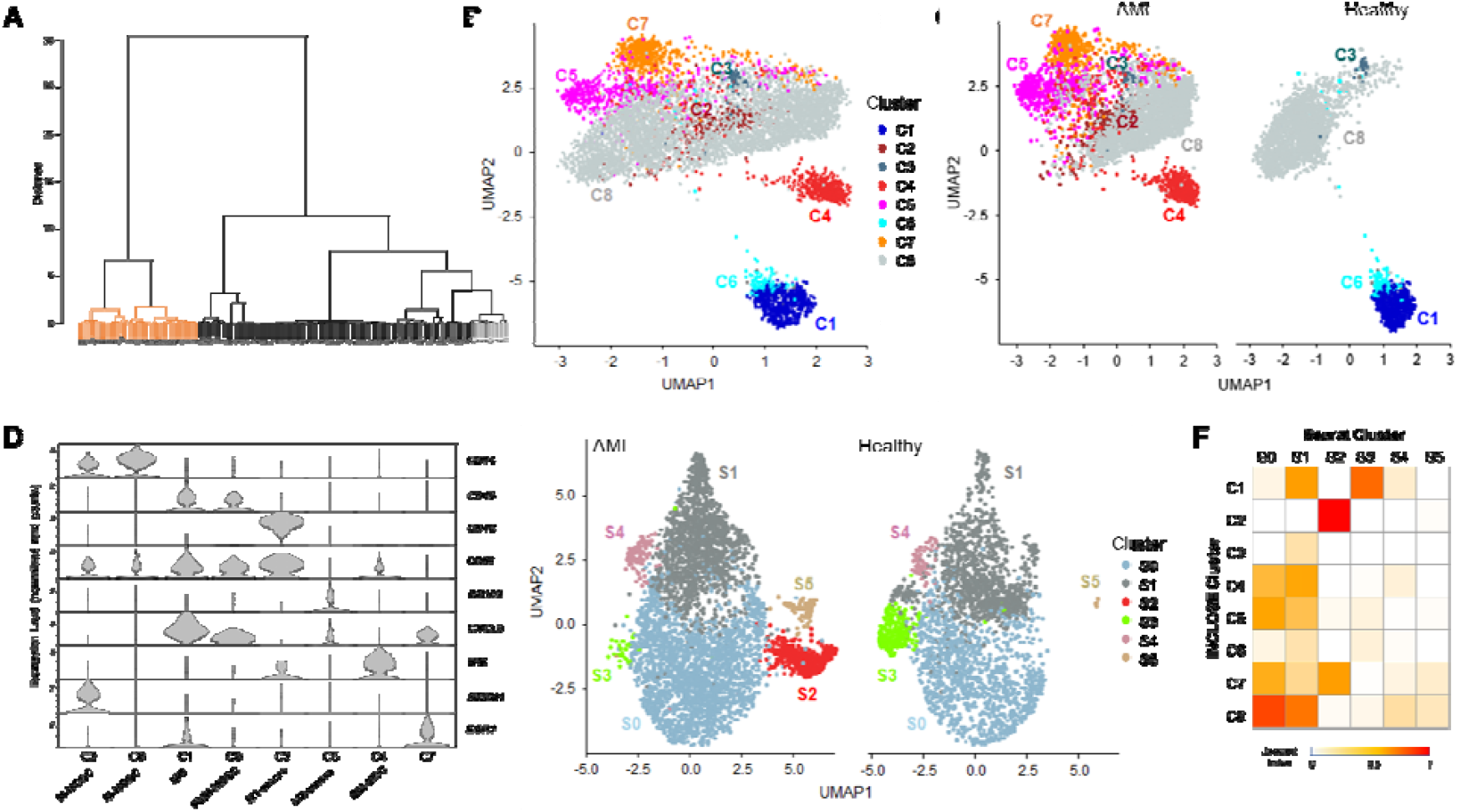
INCLOSE analysis of the MC data. (**A**) Dendrogram shows relationship between different clustering results. Orange color indicates clustering results with segregation score >0.9. (**B**) Eight clusters identified by INCLOSE are displayed in UMAP plot based on the top 2,000 variable genes. (**C**) Separate UMAP plots of cells from AML samples or frosm healthy samples show condition-specific clusters identified by INCLOSE. (**D**) Violin plots show the eight cell clusters exhibit distinct expression patterns of immune cell markers. (**E**) Six clusters identified by Seurat are displayed on the UMAP plots based on the top 10 principal components. (**F**) Bidirectional set matching based on Jaccard Index revealed correspondence between INCLOSE clusters and Seurat clusters.

We closely examined one of the clustering results where eight clusters were formed (**Fig. 2B**). Some of these clusters consisted exclusively of cells from the healthy samples or the AML samples, while others are a mixture of both (**Fig. 2C**). We compared the gene and surface protein expression levels among these clusters, which confirmed each cluster possessed a distinct molecular profile (**s**). We then proceeded with cluster annotation based on immune cell type markers ^12-14^ (**Fig. D**). The largest cluster c8, comprising a mixture of AML and healthy cells, was identified as CD14(+)CD15(-) monocytic myeloid-derived suppressor cells (M-MDSCs). Intriguingly, the AML-specific cluster c2 also exhibited the CD14(+)CD15(-) marker of M-MDSCs, but it notably expressed a leukemia marker gene *STMN1*. Cluster c5, expressing *CD68, CD163* and *MRC1* markers, was identified as M2-like macrophages, commonly known as tumor-associated macrophages (TAMs) that play a pivotal role in immune suppression ^15^. Furthermore, cluster c4, expressing *IFI6, IFNGR2*, and *HLA-DR* markers, was identified as bone marrow mesenchymal stem cell (BM-MSC), which also have immune suppression activity ^16^. The exclusive presence of clusters c4 and c5 in AML samples strongly suggested an expansion of immunosuppressive TAMs and BM-MSCs. Conversely, clusters c1 and c6, found exclusively in healthy samples, were identified as mature granulocytes (MGs) and polymorphonuclear MDSCs (PMN-MDSCs), respectively based on the CD14(-)CD15(+) marker, while cluster c6 exhibited a more pronounced inflammatory gene expression such as *CXCL8*.

For comparison, clustering using Seurat weighted nearest neighbor method ^6^ identified six clusters, with only one cluster showing specificity to the sample group (**Fig. E**). Based on the alignments between Seurat and INCLOSE clusters (**Fig. F**), this AML-specific Seurat cluster s2 corresponded to INCLOSE cluster c2 and a portion of cluster c7. No Seurat clusters matched the prominent AML-specific clusters c4 and c5 that strongly indicated immune suppression in the INCLOSE analysis. Two clusters from Seurat analysis contained predominantly but not exclusively cells from leukemia or healthy samples. However, they did not correspond to cell types associated with immune activation or suppression.

In summary, we introduced a novel INCLOSE method that integrates single-cell multi-omics data and sample metadata in cell clustering analysis. It is suited for detailed exploration of subsets of cells after the initial clustering of all cells to identify condition-specific populations. This approach complements existing methods to facilitate comprehensive single-cell analysis. INCLOSE is publicly available as an R package from https://github.com/liliulab/INCLOSE.

## Online Methods

### INCLOSE algorithm

#### Definitions and notations

Let {*T*_l_, *T*_2_,…, *T*_n_J be a set of feature matrices, each representing a specific omic profile of s single cells. For example, *T*_l_ is a gene expression matrix obtained from scRNA-seq data, *T*_2_ is a surface protein abundance matrix measured by antibody derived tags (ADT), and *T*_3_ is a matrix of phenotype features (e.g., disease group, treatment group, clinical characteristics, etc.). In these feature matrices, rows correspond to cells and columns correspond to features (**Fig. 1A**).

#### Estimation of cell-cell similarities

For each feature matrix *T*_*i*_, we compute a matrix *A*_*i*_ containing pairwise correlations between cells,

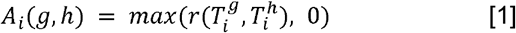

where 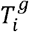 and 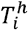 are feature vectors of cells *g* and *h*, respectively, and *r*(., .) is the correlation coefficient (Spearman or Pearson) of two feature vectors. INCLOSE supports Spearman and Pearson correlation - Spearman correlation is appropriate for studying nonlinear associations and Pearson correlation is suitable for capturing linear relationships.

We then fuse the correlation matrices in a cell-cell similarity matrix,

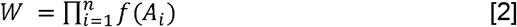

where 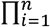 is element-wise multiplication, and *f*(.) is a Gaussian smoothing function,

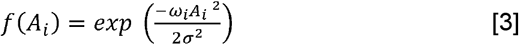

The smoothing parameter σ is shared across all data modalities to reduce the noise caused by minor variations in correlation coefficients. The weight *ω*_*i*_ is assigned to each data modality to balance the contribution to the combined similarity. *w* holds high values for pairs of cells that are similar across multiple data modalities.

INCLOSE can accommodate metadata containing a single feature, such as the sample groups each cell belongs to. In this case, the corresponding correlation matrix *A*_*i*_ is a *s* × *s* square matrix where values of cells in the same group are set to 1 and values of cells in different groups are set to 0. Next, we smooth these values using a logistic sigmoid function,

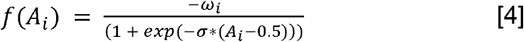

The overall similarity between a cell and all the other cells is measured as

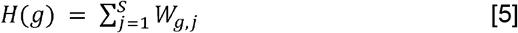

High *H* values indicate cells that share similar profiles with many other cells.

Identification of Clusters: The clustering process is similar to the one proposed by Maneck, *et al*. ^10^. It starts by identifying the cell *g*_0_ that has the highest overall similarity. Using *g*_0_ as the seed, we grow this cluster *c* by progressively adding cell *g*_k_ that maximizes the within-cluster similarity.

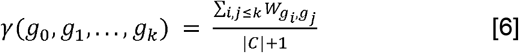

where |*c*| is the number of cells in the cluster. The iteration is terminated if the within-cluster similarity stops increasing. Among the remaining cells, we repeat the steps of selecting the seed cell and growing the cluster until no cluster with more than 5 cells could be formed or the user-specified maximum number of clusters is reached. If more than 1% of cells remain unclustered at the end of the iteration, they are assigned to a “trash cluster”.

#### Parameter tuning and optimization

INCLOSE uses the tuning parameter *ω*_*i*_ to balance the influence of feature matrix *T*_*i*_ on the clustering results. As the *ω*_*i*_ increases from 0 to 1, the influence of the feature matrix *T*_*i*_ increases. The sum of *ω*_*i*_ values are normalized to 1. The parameter σ, which governs the extent of smoothing, impacts the size of resultant clusters. As σ value increases, the size of the clusters increases, and the number of clusters decreases. σ is constrained within the range [0,1].

INCLOSE optimizes the σ and *ω*_*i*_ values by performing a grid search with a series of *ω*_*i*_ values in the range of 0 and 1 and *σ* values in the range of 0.03 and 0.3. The goal is to find a combination of σ and *ω* values that maximizes the within-cluster similarities and between-cluster distances. Because clinical and phenotypic features have already contributed to the clustering steps, they are not used again in the parameter tuning step. As such, the parameters are tuned to fit the molecular profiles.

Given a specific combination of σ and *ω* values, INCLOSE groups the cells into a set of disjoint clusters excluding the trash cluster. Using an omic profile *T*_*i*_, the within-cluster similarity metric *ϕ*_*i*_ quantifies the average similarity between pairs of cells in the same cluster,

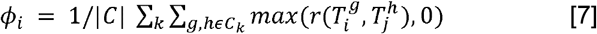

where *g* and *h* are two cells in the *k*^*th*^ cluster *c*_*k*_, *r*(.,.) is Pearson correlation coefficient, and |*c*| is the number of cells not in the “trash cluster”. The overall within-cluster similarity across multiple omes is a weighted sum

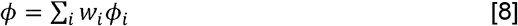

where *w*_*i*_ is the re-standardized weight computed from *ϕ*_*i*_ to ensure that *Σ w*_*i*_ =1 after excluding the metadata modality.

To calculate between-cluster distances, we first find the centroid of each cluster. For cluster *c*_*k*_, the centroid 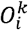 based on an omic profile *T*_*i*_ is a vector in which each element is the mean value of a feature over all cells in this cluster,

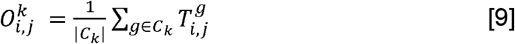

where 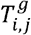 is the abundance of *j*^*th*^ feature in cell *g*, i.e., value in row *g* and column *j* in matrix *T*_*i*_, and |*c*_*k*_ | is the number of cells in cluster *c*_*k*_, i.e., the cluster size. The distance between two clusters *c*_*u*_ and *c*_*v*_ is

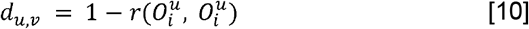

and *r*(.,.) is the Pearson correlation coefficient. We then derive the mean between-cluster distance over all clusters as

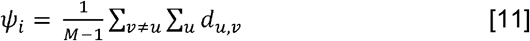

where *M* is the number of non-trash clusters. The overall between-cluster distance across multiple omes is a weighted sum

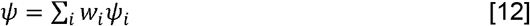

The within-cluster similarity and between-cluster distance are combined to produce a segregation score,

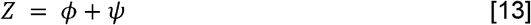

We calculate *Z* for clustering results using each unique combination of σ and ω values. A high *Z* score indicates a good clustering result, in which cells within the same cluster are highly similar and cells in different clusters are highly dissimilar. Users can use the *Z* score to guide the selection of the final clustering results.

#### Visualization of clustering results

The segregation *z* score is a heuristic measure of the quality of clustering results. In practice the best *z* score does not necessarily indicate the most biologically meaningful clustering result. Furthermore, while some parameter combinations produce very different clustering patterns, other combinations may cause little changes of the clustering patterns. We therefore provide functions to visualize and align the large number of clustering results produced by INCLOSE. To measure the similarity between two clustering results, we calculate the J-score ^11^ that aligns clusters from two clustering based on cells that are consistently grouped together (i.e., mutual presence). J-score of 1 indicates that the two clustering results are identical. Low J-score indicates the two clustering results are highly discordant. After computing J-scores for each pair of clustering results, we perform hierarchical clustering analysis to group these results into a tree structure. A clade in this tree consists of a collection of clustering results that are similar to each other. Users can choose a node and display a UMAP graph to visualize the clustering result.

### CITE-seq Data and Initial clustering using Seurat

#### Data preprocessing

The CITE-seq data of the AML study was downloaded from NCBI GEO database (accession number: GSE220473). Bone marrow tissues of three age-matched normal donors and eight individuals newly diagnosed with AML were collected, including 20,740 cells from the normal group and 20,987 cells from the AML group. For each cell, the transcriptome profile and a panel of 131 ADTs tagging cell surface proteins were measured. The raw CITE-seq count matrices were loaded into R (v4.0.3) and processed using the Seurat R package (v4.1.2). Cells with less than 100 detected genes and genes detected in fewer than 5 cells were filtered out. Cells with mitochondrial gene expression greater than 5% of the total gene expression were also removed. The RNA expression levels were normalized using standard normalization to correct for batch effects. The protein expression levels represented by ADTs, were normalized using centered log ratio normalization and scaling. A Seurat object was constructed for both the scRNA and ADT data, and the two objects were integrated using the Seurat integration pipeline.

#### Initial clustering of all cells using Seurat

We applied the Seurat FindVariableFeatures function to retrieve the top 2,000 highly variable genes and 15 highly variable ADTs across all cells. We then performed principal component analysis (PCA) analysis of these highly variable features. Based on the top 30 principal components (PCs), we used the Seurat weighted nearest neighbor pipeline to cluster the cells. At the resolution of 0.8, Seurat reported 10 distinct clusters each containing at least five cells. To identify marker genes for each cluster, we employed the FindAllMarkersMAESTRO function from the MAESTRO package ^7^. We then annotated the clusters using RNAAnnotateCelltype function based on canonical marker genes for immune cell types provided by Azimuth ^6^. The distinct cell types identified were CD14+ monocyte, CD16+ monocyte, hematopoietic stem and progenitor cell, erythroid cell, CD4 central memory cell, CD4 naïve T cell, natural killer cell, CD8 effector memory T cell, naïve B cell, and plasma blast cell. We searched for myeloid cells (MCs) in the clusters representing CD14+ and CD16+ monocyte populations by applying a gating strategy on ADTs that requires expression of CD11b(+), CD33(+), and HLA-DR (low) surface protein markers ^8^. We identified a total of 6,777 MCs including 2,543 cells in the control samples and 4,234 cells in the AML samples.

### Secondary clustering of MCs

#### Secondary clustering using Seurat

We used the same Seurat pipeline as in the initial clustering analysis but focused on MCs. We retrieved 2,000 highly variables and 15 highly variable ADTs among MCs and set the clustering resolution of 0.15. This secondary clustering analysis by Seurat revealed distinct subpopulations among MCs.

#### Secondary clustering using INCLOSE

Using the highly variable genes and ADTs among MCs identified above, we created an RNA expression matrix and an ADT expression matrix, respectively. The metadata matrix contains a single column indicating sample group (0 for healthy and 1 for AML). We explored a series of tuning parameters including σ ranging from 1/3 to 0.1/3, weights assigned to the RNA and ADT feature matrix (ω_g_ and ω_p_, respectively) ranging from 0.1 to 0.5, and weight assigned to the metadata matrix fixed at 0.03. A total of 177 unique combinations of these tuning parameters were explored, producing 177 clustering results.

#### Aligning Seurat and INCLOSE clusters

We performed bidirectional set matching between Seurat clusters and INCLOSE clusters using the J-score package ^11^. Given a Seurat cluster, Jaccard index (JI) was computed between this cluster and each INCLOSE cluster. The INCLOSE cluster showing the highest JI score was considered best match. Similarly, the Seurat cluster best matched to an INCLOSE cluster was identified. A heatmap based on JIs was created to help visualize the alignment between Seurat and INCLOSE clustering results.

