## Supplemental Figures 1-4 for "Discovering Condition-specific Cell Populations via Integrative Clustering of Single-cell Data"

A

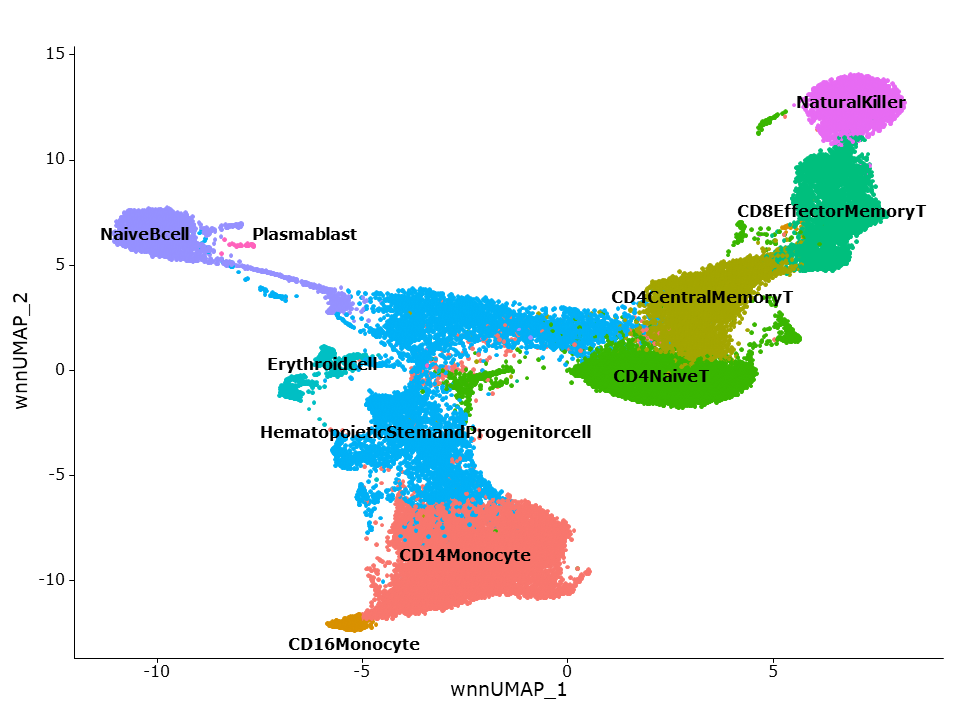

B
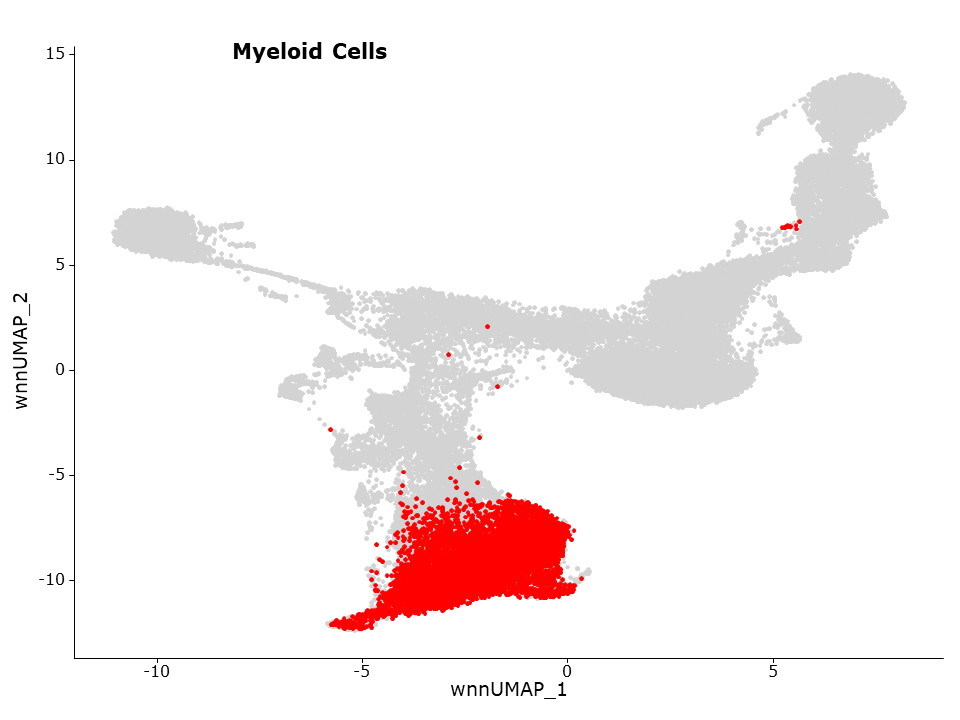

**Supplementary Figure 1**. Cell clusters identified by Seurat analysis of the entire AML data set. (**A**) UMAP plot shows clusters of cells with annotated cell types. (**B**) UMAP plot highlights the myeloid cells that will be further clustered by INCLOSE to discover subpopulations.

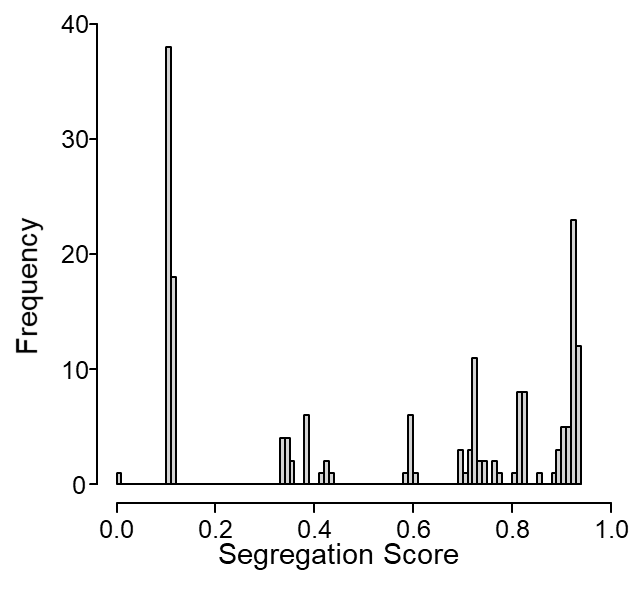

**Supplementary Figure 2**. Histogram shows distribution of segregation scores of 177 clustering results using various combinations of tuning parameter values.

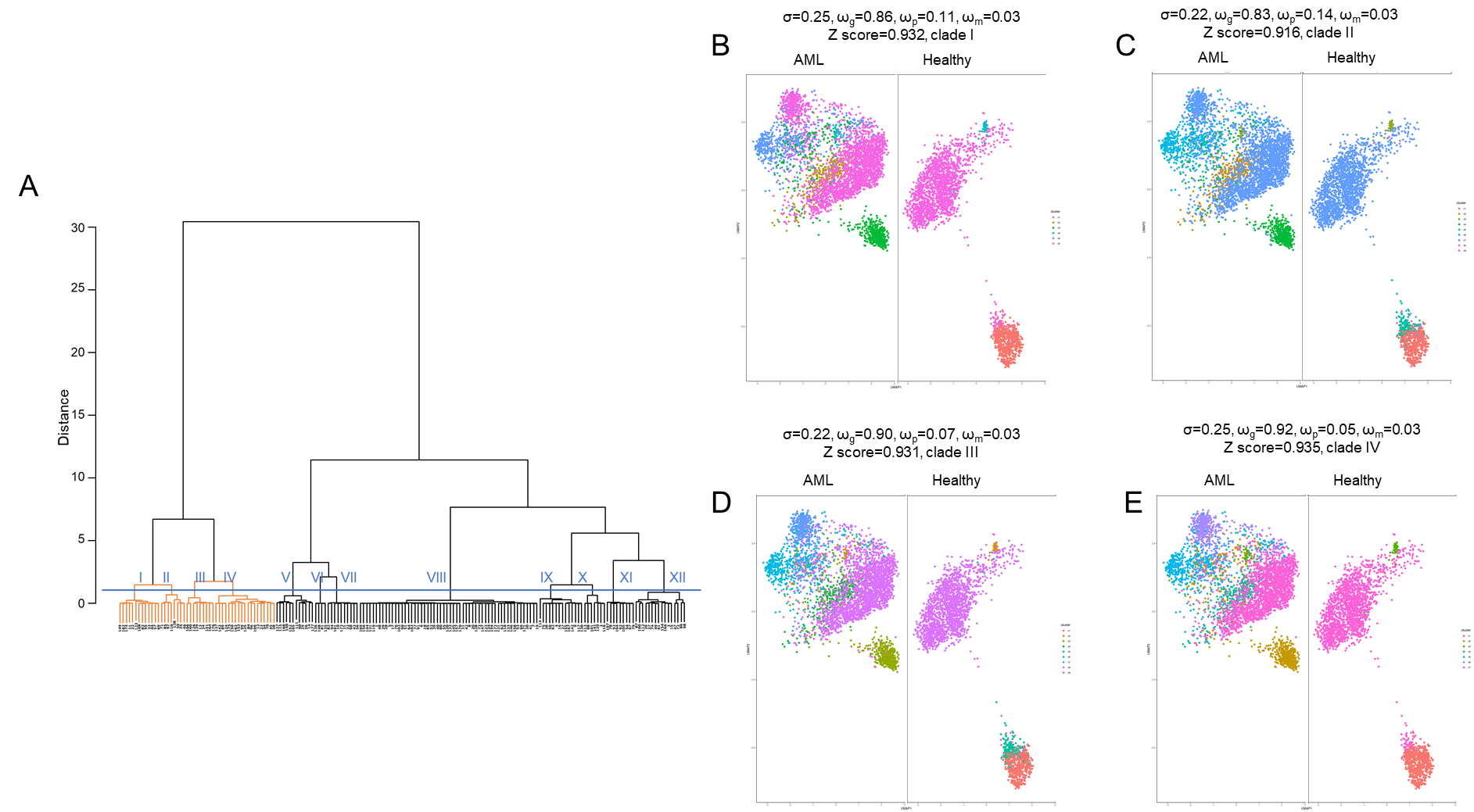

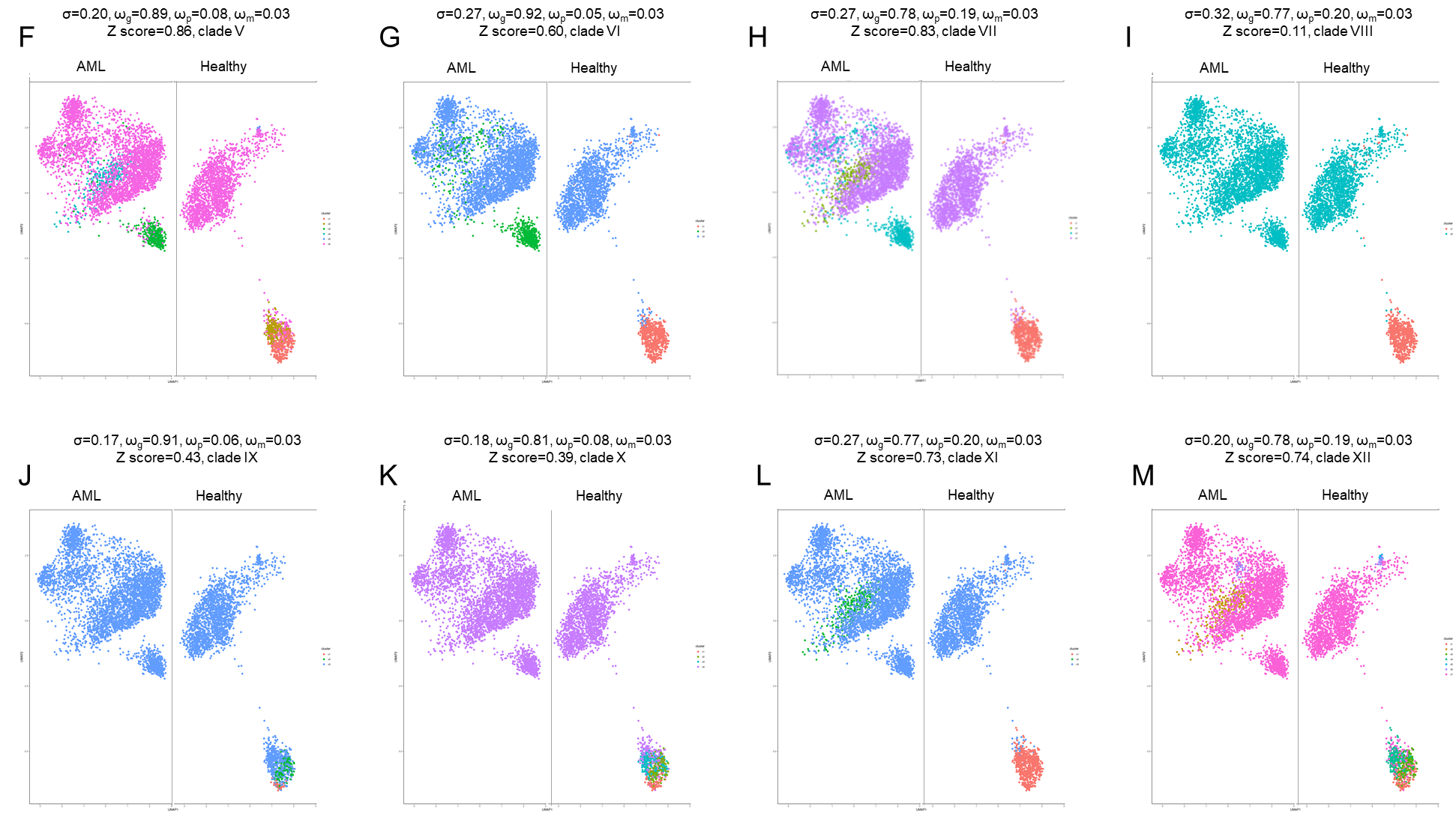

**Supplementary Figure 3**. INCLOSE clustering results using different tuning parameters. (**A**) Dendrogram of the 177 clustering results. Orange color indicates clustering results with segregation score >0.9. The horizontal blue line cuts the dendrogram into twelve clades (I – XII). (**B-M**) In each clade, the clustering result with the highest segregation score was plotted in UMAP. Values of the tuning parameters are displayed, including smoothing parameter (σ) and weights for gene expression (ω_g_), surface protein expression (ω_p_), and metadata (ω_m_). Different colors represent different cell clusters.

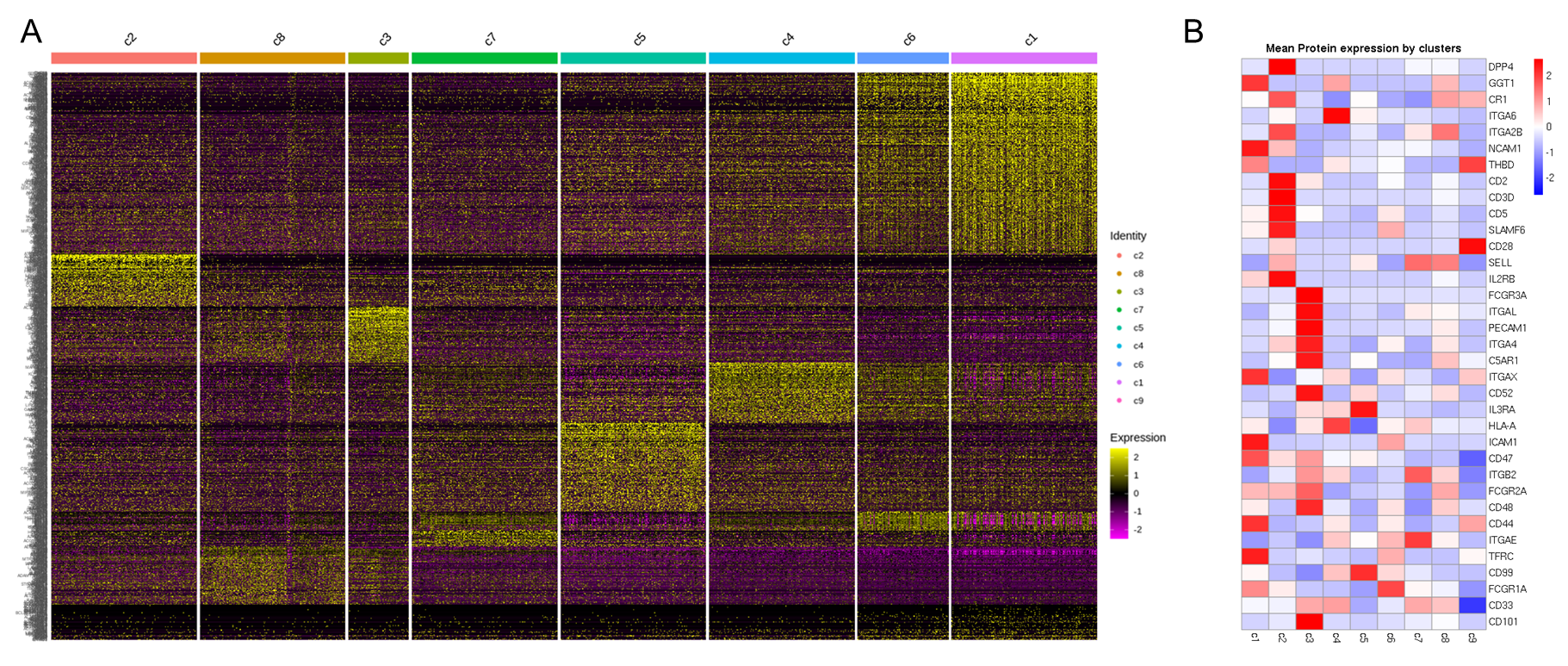

**Supplementary Figure 4**. Distinct molecular profiles of the eight clusters identified by INCLOSE analysis. Heatmaps shows differentially expressed genes (**A**) and differentially expressed surface proteins (**B**) between clusters. Comparison was done between a given cluster and the remaining clusters.
